# Effects of food availability on larval development during ontogenetic niche shift in a marine annelid

**DOI:** 10.1101/2025.04.20.649073

**Authors:** Nadine Randel

## Abstract

Many marine invertebrates have a biphasic life cycle with a free-swimming larva and a bottom-dwelling adult. The transition from a planktonic to a benthic lifestyle is a significant step in the animal’s life history, highly regulated and influenced by external and internal factors. Since the readiness to settle and the presence of a suitable seafloor habitat do not always coincide, larvae sometimes need to extend their planktonic phase. Little is currently known regarding how larvae partition their energy for coordinating development and growth according to food type and availability in their settlement habitat. Here, I investigate the effect of food availability and type on development in *Platynereis dumerilii* larvae. I assessed cell proliferation, growth, and feeding onset over six days using two different food sources. The results indicate that food availability and type affect larval growth, with starved larvae slowing development and conserving resources, whereas fed larvae allocate resources to brain development and posterior growth. Overall, this work contributes to our understanding of how competent marine larvae regulate the duration of their planktonic phase and how nutritional status affects development.

## Introduction

Every animal is adapted to its ecological niche and reacts to its environment. However, many animals change their niche during their lifetime. This can be as simple as an increase in body size, necessitating different food sources and shelters [1]. The transition between different niches and the resulting reorganisation of functional systems (e.g., nervous system, and locomotor system) has been termed ontogenetic niche shift [2]. Ontogenetic niche shift occurs in many invertebrate and vertebrate taxa such as insects, molluscs, annelids, echinoderms, fishes, and amphibians [3,4]. It is an important step in an animal’s lifetime, essential for gaining sexual maturity and preserving ecological communities. In particular, the transition of planktonic marine larvae to bottom-dwelling adults has recently come into focus due to its significance for marine ecosystems, fisheries, aquaculture, and biofouling [5–8].

A hallmark of these marine invertebrate taxa is the biphasic life cycle. The free-swimming planktonic larvae inhabit the water column from a few hours to several months before they settle down to the seafloor [9]. Feeding larvae normally spend longer in the plankton, whereas a short planktonic phase is often associated with lecithotrophy, where larvae obtain their nourishment from maternally provided yolk or lipid droplets.

By the end of the larval stage, the larva has developed competency, the ability to respond to environmental cues and settle onto the seafloor [10–15].

Due to the sometimes highly specific nature of environmental cues required to induce settlement, competent larvae might not be able to settle. As a result, they either spontaneously metamorphose, die within a few days, or survive an extended period in the plankton but lose the ability to develop into adults [9]. Consequently, the ability to maintain competency is important for the animal’s fitness.

Prior studies have shown that starved planktotrophic larvae can prolong their larval phase [16–18]. However, little is known about the ability to maintain competency in planktotrophic and lecithotrophic larvae [16]. Additionally, lecithotrophic larvae may be incapable of feeding, while others are facultative planktotrophic, or, as in the marine annelid *Platynereis dumerilii*, they settle as pre-metamorphic hemisessile feeding larvae [19–21].

Despite the importance of maintaining larval competency, our understanding of the underlying mechanisms remain limited. Although prior studies have highlighted the role of food in many species, its effects on development remain poorly understood. This study investigates how food availability affects the development of competent *P. dumerilii* larvae. *Platynereis* is a well-established model with a biphasic life cycle, featuring a lecithotrophic larva that develops into a bottom-dwelling adult (Figure 1A) [19]. After hatching, the gut of the larva is not fully formed, and four maternally provided lipid droplets serve as an initial energy source. Later they feed on various species of green algae and diatoms on the seafloor, including the microalgal species used in this study: the unicellular, free-swimming green alga *Tetraselmis suecica*, and the benthic, biofilm-forming diatom *Grammatophora marina*.

**Figure 1:**
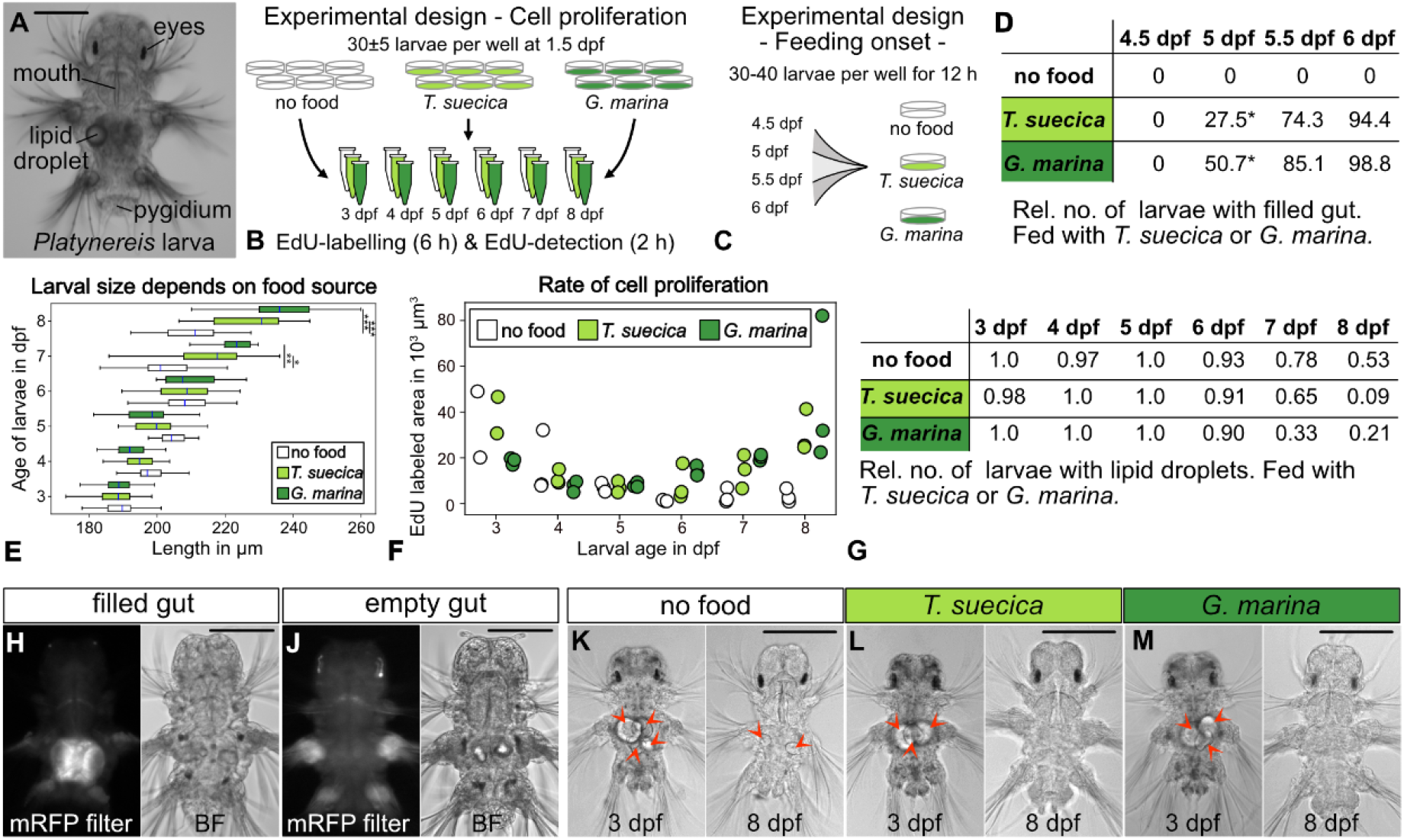
Effect of microalgae on larval size, cell proliferation, and feeding behaviour during ontogenetic niche shift. (A) Light micrograph of a 6 dpf old Platynereis larva. (B) Experimental design of the cell proliferation assay, and (C) testing of feeding onset. (D) Proportion of larvae with microalgae in the gut (*p ≤0.05). (E) Box and whisker plots of larval size across ages and food conditions (p-values: * ≤0.05, **≤0.001, ***≤0.0001). (F) Total area of EdU-labeled cells from three larvae per age and food conditions. (G) Proportion of larvae with 1-4 lipid droplets, feeding on T. suecica and G. marina. (H-J) Light and fluorescence micrograph of T. suecica fed and unfed larvae fed, imaged with RFP filter. (K-M) Light micrograph of 3 and 8 dpf old larvae with maternally provided lipid droplets (arrowheads), under different food conditions. (D,E,G) Statistics: paired t-test, Holm-Bonferroni correction. Scale bar: 100 μm.

My work provides new insights into how food availability affects the timing of feeding onset, use of the maternally provided lipid droplets, and larval growth rates.

## Material and methods

### *Platynereis dumerilii* and microalgae culture

Animals and microalgae are cultured in the Marine Invertebrate Culture Unit (MICU) at the University of Exeter, UK and were provided by the laboratory of Dr Elizabeth Williams [22].

### Diatom biofilm preparation

To enhance the biofilm formation, 6-well plates (ThermoFisher Scientific 130 184 bioLite 6-well multidish) were coated with Poly-D-lysine (Sigma-Aldrich P6407, final concentration 5 μg/ml) for 2 hours at 37ºC and subsequently dried at 50ºC. The coated 6-well plates were then incubated for 36 hours with 3 ml *Grammatophora marina* diatom culture medium. The medium consisted of filtered seawater supplemented with F/2 nutrient medium (1 ml per 2.5 l; ZM Systems), and a silicate solution (1ml per 2.5 l; ZM Systems). Afterwards, the 6-well plates were gently washed and covered with 1 μm-filtered UV-light sterilised artificial sea water (Tropic Marin Pro-Reef salt) at a salinity of 33 ppt. To ensure food was not a limiting factor, an excess was provided. The biofilm coverage was calculated using Biofilm_coverage_macro.ijm [22]. It was estimated at 6%-12% (Suppl. File 1).

### Experimental design for cell proliferation assay

30±5 larvae from two batches of larvae with differing parentage were added to the prepared 6-well plates, with three biological replicates. The 1.5 day post fertilisation (dpf) larvae were raised either on *G. marina* biofilm, or fed with 50 μl *Tetraselmis suecica* algae culture. The larvae were kept in a 18ºC incubator with 18 h light : 6 h dark photoperiod.

### Cell proliferation assay and DAPI staining

Newly synthesised DNA was labelled in whole larvae and detected using the Click-IT EdU Alexa Fluor 647 Imaging Kit (Invitrogen C10340) according to the kit instructions. *Platynereis* larvae were incubated with EDU at 3, 4, 5, 6, 7, and 8 dpf for 6 h. Afterwards, the larvae were fixed for 1 h with 4% PFA (Agar Scientific AGR1026) in PBW (phosphate buffer [PBS] + 0.1% Tween20 [Sigma-Aldrich P9416]) and washed in PBW. After a final washing step with PBS, the larvae were incubated for 2 h with the reaction cocktail (EdU kit) and washed in PBS. Finally, larvae were stained with DAPI (0.5μmol/ml) (Merck D9542-1mg) for 10 min, and stored in 2,2 Thiodiethanol (Merck 166782-500g) at 4ºC in the dark until imaging. Sample size per age and condition is 12-51 larvae (Figure 1B).

### Experimental design for feeding onset characterization

Thirty to forty larvae from two batches were incubated in 6-well plates with *G. marina* biofilm, an excess of *T. suecica* algae culture, or no food (three biological replicates) at 4.5, 5, 5.5, and 6 dpf for 12 h (Figure 1C). After fixation with 4% PFA in PBS for 1 h, the larvae were washed with PBS and imaged within 2 weeks.

### Imaging

#### Confocal microscopy imaging of EdU-labelled larvae

Larvae were mounted in 2,2 Thiodiethanol and imaged with a Leica upright SP5 confocal microscope (40x oil objective, NA 1.25) in the imaging facility at University of Cambridge (UK), Department of Zoology.

#### Bright field microscopy imaging of larval size, lipid droplets, and gut content

Larvae were imaged with Axioscope 40 (20x air objective) and recorded with Micromanager 2.0. in the imaging facility at the University of Cambridge (UK), Department of Zoology. The microalgae’s autofluorescence was recorded with 20% light intensity and 200 milliseconds exposure time, using an RFP filter.

Length of the EdU labelled larvae was measured from the head to the pygidium using Fiji [23]. In addition, the presence or absence of maternally provided lipid droplets was recorded for each larva. According to the experimental setup, incubation time with food in the EdU labelled larvae is 36h-156h.

### Data analysis

Data analysis and plotting of larval size, lipid droplets, and gut content were performed using Python 3.7, and estimation of EdU labeled area was done using ImageJ Macros. All scripts are available on https://github.com/nrandel/Randel_2025.

#### Estimation of EdU labelled area

Each 3D image stack was converted into a 3D mask after background subtraction, using ImageJ/Fiji. Using ROIs of the head, first and second segment, and third segment and pygidium, the signal in the selected areas was extracted. Due to sample movement during imaging, the signal for one sample from 3 dpf (no food) and one from 3 dpf (*T. suecica*) could not be extracted.

#### Manual detection of EdU labelled cells

3dpf EdU-labelled larva was manually segmented in napari [24].

#### Statistical analysis of larval size

Variation between the biological replicates and normal distribution was tested with the Shapiro-Wilk test and one-way ANOVA, respectively. H_0_ was rejected for three of 51 replicates, and a parametric test was used subsequently.

Comparison between groups and age has been done with one-way ANOVA, and Holm-Bonferroni correction for multiple-comparison analysis. Pairwise analysis was performed with a paired t-test and corrected using Holm-Bonferroni correction.

#### Statistical analysis of feeding on-set and lipid droplets

Fisher’s exact test was used to test for variation between the biological replicates and a significant association between the variables. Holm-Bonferroni correction was performed for multiple-comparison analysis.

## Results

### Feeding source affects feeding onset, growth and lipid droplet consumption

To determine when the larvae start feeding and whether they have any food preferences, *P. dumerilii* was exposed to either *T. suecica* or *G. marina* for 12 hours (Figure 1C). For both algae, larvae started feeding between 5 dpf and 5.5 dpf, though significantly more larvae had fed on *G. marina* by the end of the 12 hour feeding period (Figure 1D, H-J).

Next, I assessed the growth rate and lipid droplet depletion to evaluate the effect of food on larval growth and the consumption of maternally provided resources. For this experiment, 1.5 dpf larvae were continuously raised with either *T. suecica* or *G. marina*, and their size and absence of lipid droplets were recorded (Figure 1E, G). Before feeding onset, the unfed and fed larvae grew at the same rate. However, after 7 dpf, unfed larvae grew significantly slower than fed larvae, despite still having access to the maternally provided lipid droplets (Figure 1G). After 7 dpf, the unfed larvae appeared to consume the lipid droplets at a slower rate than fed larvae, though this effect was not significant (Suppl. File 2).

### Increased cell proliferation in head and posterior growth zone

*Platynereis dumerilii* grows continuously throughout its life by adding new segments at the posterior growth zone. To investigate the number and location of new cells, I performed a cell proliferation assay for six hours each day in 3 to 8 dpf larvae (Figure 1B).

Differences in growth rate between fed and unfed larvae are reflected in the amount and timing of cell proliferation. Until 5 dpf, unfed and fed larvae develop at a similar rate, with exceptionally high cell proliferation rate in the 3 dpf larvae (Figure 1E,F). At this stage, the larva consists of over 9000 cells [25]. By manually segmenting the EdU labelled cells in one sample, I found that between 72 hpf and 78 hpf, more than 800 new cells developed (Movie 1).

However, in larvae older than 5 dpf, cell proliferation significantly differs between fed and unfed larvae. Unfed *P. dumerilii* further reduced cell proliferation throughout the body, whereas fed larvae showed increased cell division (Figure 1F). The slightly earlier onset of cell proliferation in the posterior growth zone in 6 dpf old *G. marina* fed larvae further supports the observed trend that larvae fed with the diatom grow initially faster than those fed with *T. suecica* (Figure 1F, 2K, Q). In addition, fed larvae had weak cell proliferation in the trunk segments but a noticeable increase in newly developed cells in the head, likely contributing to brain development (Figure 2).

**Figure 2:**
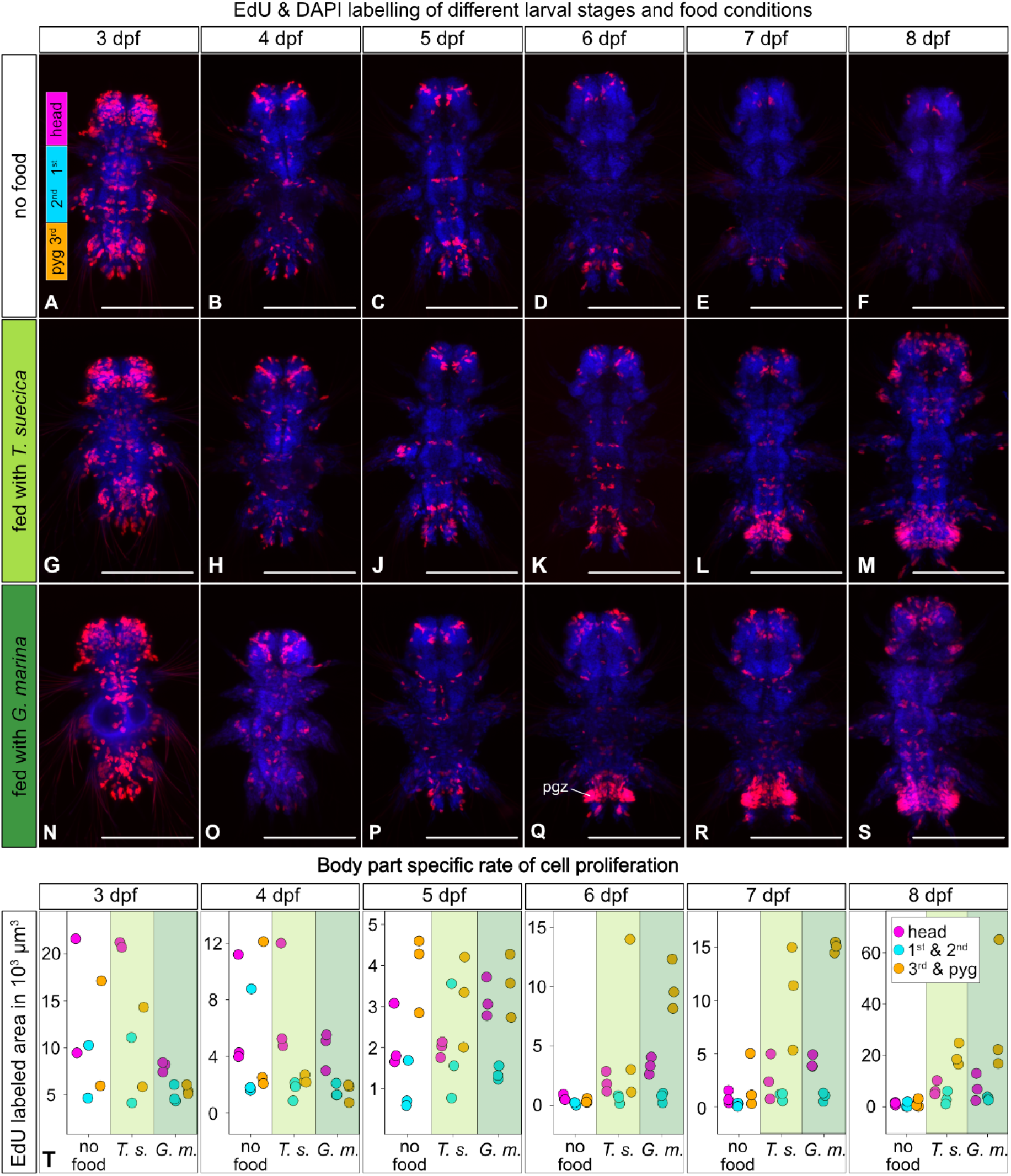
Cell proliferation across larval stages and food conditions. (A-S) Fluorescence micrograph of cell proliferation (red) counterstained with DAPI (blue). The labeled regions in (A) correspond to (T) where cell proliferation is estimated in the head, first and second segments, and third segment and pygidium. Three larvae aged three to eight days were analysed for different food conditions. pgz - posterior growth zone, G. m. - *G. marina*, T. s. - *T. suecica*. Scale bar: 100 μm.

## Discussion

In this study, I investigated how different food types affect late larval development in *P. dumerilii*. The larva can settle between three to four days post fertilization in response to the settlement cue, *G. marina* [22]. However, in the absence of *G. marina* or other food sources, such as *T. suecica*, the competent larvae delay settlement.

### Food availability affects larval development

This work demonstrates how competent *P. dumerilii* larvae respond to the lack of food by pausing cell proliferation throughout the whole body. These results complement a previous study showing that the larvae can survive for up to 30 days without food, remaining at the three-segment stage [26]. Nutritional modulation of development is not unique to *P. dumerilii*, and has been described in many other taxa, in which insufficient nutrients impact developmental trajectories, alter cell metabolism, and influence resource allocation during organogenesis [27].

Before the onset of feeding, food availability does not significantly impact growth rate and cell proliferation in *P. dumerilii*. However, once larvae begin feeding, cell proliferation is initiated in the head and posterior growth zone. In addition, the data suggest that feeding larvae may deplete the maternally provided lipid droplets faster than starved ones.

A possible explanation for these differences in resource utilization and development is that starved larvae limit energy consumption to prolong their planktonic phase, thereby increasing the time available to them for finding a suitable habitat before depleting energy resources. In contrast, feeding larvae maximize resource use of food and lipid droplets to increase their body size, which may enhance their ability to compete with conspecifics for food and space.

### Feeding onset depends on food type

My data show that *Platynereis* begins feeding between 5 and 5.5 dpf, nearly two days after reaching competency for settlement. The slightly earlier feeding onset with *G. marina* may be due to greater accessibility of the biofilm, compared to the free-swimming *T. suecica*. Alternatively, since *G. marina* induces earlier settlement, it may also accelerate larval development, enabling earlier food uptake, though evidence remains lacking [22].

### Summary and Outlook

This work paves the way for future studies on the molecular mechanisms underlying the nutritional modulation of development. Due to its small size and life history, *Platynereis* provides a valuable model for investigating the cross-talk between nutrition, metabolism, and development to gain a detailed mechanistic understanding of resource allocation, storage, and the effect of maternally provided yolk on early development.

## Data accessibility

Raw data, Python 3.7 scripts, and ImageJ Macros are available on https://github.com/nrandel/Randel_2025.

## Declaration of AI use

ChatGPT-3.5 (OpenAI 2024) has been used to support writing the scripts for data analysis and plotting.

## Author contributions

NR designed and performed all experiments, conducted data analysis, and wrote the manuscript.

## Acknowledgements

I thank Dr. Elizabeth Williams from the University of Exeter for hosting me and sharing her expertise on microalgal biofilms, as well as Dr. Susanne Vogeler and Callum Teeling for preparing the biofilm and testing the lifetime of algae autofluorescence, respectively. I am also thankful to Prof. Howard Baylis at the University of Cambridge for his support and for providing essential reagents. Thanks also to Prof. Matthias Landgraf for granting me access to the light microscopy facility at the University of Cambridge (Department of Zoology) and to Dr. Matthew Wayland for his knowledge and advice on confocal microscopy and image analysis. I would also like to thank Prof. Gáspár Jékely (COS, University of Heidelberg) and Dr. James Herbert-Read (University of Cambridge) for their advice on the manuscript.

## Footnotes

### Supplementary Files

Suppl File 1: Estimated Biofilm coverage: EdU experiment and Feeding experiment

Suppl File 2: Raw data for larval size, lipid droplets, gut content

Movie 1 - Manually segmented cells in a 78 hpf old larva

